# Cue-induced effects on decision-making distinguish subjects with gambling disorder from healthy controls

**DOI:** 10.1101/564781

**Authors:** Alexander Genauck, Milan Andrejevic, Katharina Brehm, Caroline Matthis, Andreas Heinz, André Weinreich, Norbert Kathmann, Nina Romanczuk-Seiferth

## Abstract

While an increased impact of cues on decision-making has been associated with substance dependence, it is yet unclear whether this is also a phenotype of non-substance related addictive disorders, such as gambling disorder (GD). To better understand the basic mechanisms of impaired decision-making in addiction, we investigated whether cue-induced changes in decision-making could distinguish GD from healthy control (HC) subjects. We expected that cue-induced changes in gamble acceptance and specifically in loss aversion would distinguish GD from HC subjects.

30 GD subjects and 30 matched HC subjects completed a mixed gambles task where gambling and other emotional cues were shown in the background. We used machine learning to carve out the importance of cue-dependency of decision-making and of loss aversion for distinguishing GD from HC subjects.

Cross-validated classification yielded an area under the receiver operating curve (AUC-ROC) of 68.9% (p=0.002). Applying the classifier to an independent sample yielded an AUC-ROC of 65.0% (p=0.047). As expected, the classifier used cue-induced changes in gamble acceptance to distinguish GD from HC. Especially increased gambling during the presentation of gambling cues characterized GD subjects. However, cue-induced changes in loss aversion were irrelevant for distinguishing GD from HC subjects. To our knowledge, this is the first study to investigate the classificatory power of addiction-relevant behavioral task parameters when distinguishing GD from HC subjects. The results indicate that cue-induced changes in decision-making are a characteristic feature of addictive disorders, independent of a substance of abuse.

**Remarks:** To ensure a more convenient reviewing process, we positioned figures and tables at their destined position.

## INTRODUCTION

Gambling disorder (GD) is characterized by continued gambling for money despite severe negative consequences^1^. Burdens of GD include financial ruin, loss of social structures, as well as development of psychiatric comorbidities^2^. In line with this clinical picture of impaired decision making, GD subjects have also displayed impaired decision making in laboratory experiments^3,4^.

Besides impaired decision making, cue reactivity has been a crucial concept in understanding addictive disorders including GD^5,6^. Through Pavlovian conditioning, any neutral stimulus can become a conditioned stimulus (i.e. a cue) if it has been paired with the effects of the addictive behavior^7^. In addictive disorders, including GD, cues may induce attentional bias, arousal, and craving for the addictive behavior in periods of abstinence^8,9^. Treatment of addictive disorders may focus on identifying and coping with individual cues that induce craving for addictive behavior^10^. If we understood better how cues exert control over instrumental behavior and decision-making, we would be able to improve treatment tools and even public health policy for GD and perhaps other addictive disorders. In the present study we were thus interested in broadening our understanding of the basic mechanisms of impaired decision making in addictions, especially with respect to cue-induced effects on value-based decision making. The effect of cues exhibiting a facilitating or inhibiting influence on instrumental behavior and decision making is known as Pavlovian-to-Instrumental Transfer (PIT)^11^. PIT experiments usually have three phases: a first phase where subjects learn an instrumental behavior to gain rewards or avoid punishments, a second phase where subjects learn about the value of arbitrary stimuli through classical conditioning, and a third phase (the PIT phase), where subjects are supposed to perform the instrumental task, while stimuli from the second phase (changing from trial to trial) are presented in the background. The PIT phase measures the effect of value-charged cues on instrumental behavior despite the fact that the background cues have no objective relation to the instrumental task in the foreground. For instance, certain cues could increase the likelihood of gamble acceptance or the sensitivity to the gain offered in the gamble. In the current study we focus only on the PIT phase. PIT has recently drawn attention in the study of substance use disorders (SUDs)^12^. This is because PIT effects can persist even when the outcome of the instrumental behavior has been devalued^13^, and because increased PIT has been associated with a marker for impulsivity^14^ and with decreased model-based behavior^15^. Lastly, PIT effects tend to be stronger in subjects with a substance-use-disorder than in healthy subjects^12,16^, and increased PIT has been associated with the probability of relapse^12^.

Increased PIT effects are based on Pavlovian and instrumental conditioning and on their interaction. This highlights how addictive disorders rely on learning mechanisms^17^. GD is an addictive disorder independent of any influence of a neurotropic substance of abuse. The study of PIT in GD may thus further shed light on whether increased PIT in addictive disorders is a result of learning, independent of any substance of abuse, or even a congenital vulnerability^18^. We are aware of three studies that have observed in GD subjects increased cue-induced effects on decision-making and instrumental behavior, comparable to increased PIT effects. In two single-group studies, GD subjects have shown higher delay discounting (preferring immediate rewards over rewards in the future) in response to a casino environment vs. a laboratory environment^19^ and to high-craving vs. low-craving gambling cues^20^. In a third study, GD subjects have been more influenced than HC subjects by gambling stimuli in a response inhibition task^21^. To our knowledge, however, there are no studies yet that have investigated the effect of cue reactivity on loss aversion in GD as a possibly relevant PIT effect in GD.

Loss aversion (LA) is, besides delay discounting, another facet of value-based decision-making. It is the phenomenon wherein people assign a greater value to potential losses than to an equal amount of possible gains^22^. For example, healthy subjects tend to agree to a coin toss gamble (win/loss probability of 0.5) only if the amount of possible gain is at least twice the amount of possible loss. In GD subjects, LA seems to be reduced^23,24^, but there are also studies that have found no difference in LA between GD and HC subjects^25^.

High LA protects against disadvantageous gambling decisions. However, it has been observed that LA can be transiently modulated by experimentally controlled cues^26^ and that this LA modulation varies considerably across subjects^27^. In GD subjects, loss aversion might be particularly cue-dependent leading to reckless gambling especially in casino contexts or at slot machines. In the current study, we thus hypothesized that GD subjects should show stronger PIT effects than HC subjects in their gambling decisions and especially stronger drops in LA when e.g. gambling-related cues are present (i.e. higher “loss aversion PIT”).

So far, we have mentioned studies that have used group-mean difference analyses to investigate decision making or cue reactivity in addictive disorders. This approach is faithful to the desire to explain human behavior rather than predict it^28^. However, this may lead to overly complicated (i.e. overfitted) models, which do not correctly predict human behavior in new samples^28^. Thus, in the current study we wanted to avoid overfitting and isolate a model with not only explanatory but also predictive value^28^. We did so by disentangling the specific benefits of “loss aversion PIT” parameters when distinguishing GD from HC subjects. Hence, we used machine learning methods in addition to classical mean-difference statistics to test our hypotheses. This approach has drawn increasing attention in the field of clinical psychology and psychiatry^29^. In particular, we built and tested an algorithm that decides between various loss aversion models and different models with and without PIT to classify subjects into HC vs. GD groups. Importantly, to avoid overfitting, we used out-of-sample classification^30–32^. Our results allowed us to disentangle which PIT effects are relevant to distinguish GD from HC subjects.

When selecting cues for this study, we aimed at expanding on existing studies investigating cue-effects in GD^19–21^. Besides gambling-related cues, we thus selected additional cues from different motivational and emotional categories^12^ related to GD. These categories comprised images used in gambling advertisements as well as for advertisement of GD therapy and prevention (positive and negative cues).

We expected that our classifier would select models that incorporate the modulation of loss aversion by gambling and other emotional cues (“loss aversion PIT”) to distinguish between HC and GD subjects.

## MATERIALS AND METHODS

### Samples

GD subjects were diagnosed using the German short questionnaire for gambling behavior questionnaire (KFG)^33^. The KFG diagnoses subjects according to DSM-IV criteria for pathological gambling. Scoring 16 points and over means “likely suffering from pathological gambling”. However, here we use the DSM-5 term “gambling disorder” interchangeably, because the DSM-IV and DSM-5 criteria largely overlap. The GD group were active gamblers and not in therapy. The HC group consisted of subjects that had no to little experience with gambling, reflecting the healthy general population as in other addiction studies^5^. For further information on the sample, see **Tab. 1** and **Supplements (1.1)**. GD and HC were matched on relevant variables (education, net personal income, age, alcohol use), except for smoking severity. We thus included smoking severity in the classifier and tested it against classifying based only on smoking severity. For final validation of the fitted classifier we used a sample from another study where subjects performed the affective mixed gambles task in a functional magnetic resonance imaging (fMRI) scanner (see **Tab. S2**)^34^.

### Procedure and data acquisition

Subjects completed the task at the General Psychology behavioral lab of the Department of Psychology of Humboldt-Universität zu Berlin. They were sitting upright in front of a computer screen using their dominant hand’s fingers to indicate choices on a keyboard. Subjects were attached five passive facial electrodes, two above musculus corrugator, two above musculus zygomaticus, and one on the upper forehead. We recorded electrodermal activity (EDA) from the non-dominant hand. Subjects of the validation sample completed the task in an fMRI environment (head-first supine in a 3-Tesla SIEMENS Trio MRI at the BCAN - Berlin Center of Advanced Neuroimaging). Results of the fMRI and peripheral-physiological recordings will be reported elsewhere.

### Affective mixed gambles task

We were inspired by established tasks to measure general LA and LA under the influence of affective cues^27,35^. Subjects were each given 20€ for wagering. On every trial, subjects saw a cue that they were instructed to memorize for a paid recognition task after the actual experiment. After 4s (jittered), a mixed gamble, involving a possible gain and a possible loss, with probability P = 0.5 each, was superimposed on the cue. Subjects had to choose how willing they were to accept the gamble (**Fig. 1A**) on a 4-point Likert-scale to ensure task engagement^35^. Subjects of an independent validation sample completed the task in an fMRI scanner and had an additional wait period to decide on the gamble (**Fig. 1B**). Gambles were created by randomly drawing with replacement from a matrix with possible gambles consisting of 12 levels of gains (14, 16, …, 36) and 12 levels of losses (−7, −8, …, −18). This matrix is apt to elicit LA in healthy subjects^23,35^. Outcomes of the gambles were never presented during the task but subjects were informed that after the experiment five of their gamble decisions with ratings of “somewhat yes” or “yes” would be randomly chosen and played for real money. As affective cues, four sets of images were assembled: 1) 67 gambling images, showing a variety of gambling scenes, and paraphernalia (*gambling cues*) 2) 31 images representing negative consequences of gambling (*negative cues*) 3) 31 images representing positive effects of abstinence from gambling (*positive cues*): 4) 24 neutral IAPS images (*neutral cues*). For further information on validation of the cue categories and on access to the stimuli, please see **Supplements (1.2)**. We presented cues of all categories in random order and each gambling cue once. For negative, positive, and neutral cue categories, we randomly drew images from each pool until we had presented 45 images of each category and each image at least once. Hence, we ran 202 trials in each subject. Gambles were matched on average across cue categories according to expected value, variance, gamble simplicity, as well as mean and variance of gain and loss, respectively. Gamble simplicity is defined as Euclidean distance from diagonal of gamble matrix (*ed*)^35^. HC showed on average missed trial, GD 1.05 (no significant group difference, F = 0.022, p = 0.882). In fMRI validation study, HC: 3.13, GD: 4.10, (no significant group difference, F = 0.557, p = 0.457).

**Figure 1:**
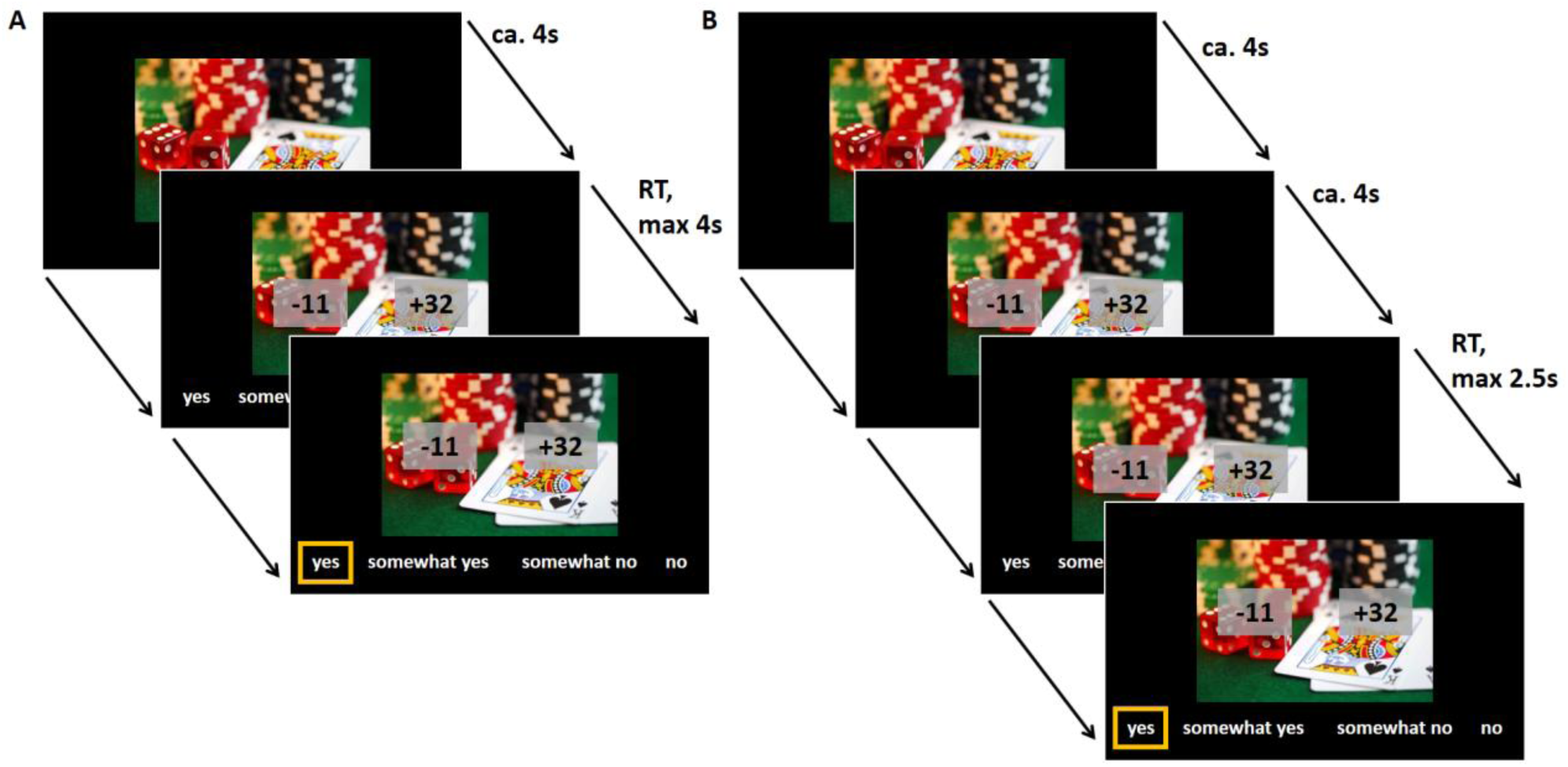
The affective mixed gambles task. One trial is depicted. **A:** behavioral sample. **B:** fMRI validation sample. Subjects first saw a fixation cross with varying inter-trial-interval (ITI, 2.5s to 5.5s, up to 8s in fMRI version; not displayed here). Subjects then saw a cue with different affective content (67 of 67 gambling related, 45 of 31 with positive consequences of abstinence, 45 of 31 with negative consequences of gambling, 45 of 24 neutral images) for about 4s. Subjects were instructed to remember the cue for a paid recognition task after all trials. Then a gamble involving a possible gain and a possible loss was superimposed on the cue. Subjects were instructed to shift their attention at this point to the proposed gamble and evaluate it. In the current example, a coin toss gamble was offered, where the subject could win 32 Euros or lose 11 Euros (50/50 probability). Position of gain and loss was counterbalanced (left/right). Gain was indicated by a ‘+’ sign and loss by a ‘-’ sign. In the behavioral sample, subjects had 4s to make a choice between four levels of acceptance (yes, somewhat yes, somewhat no, no; here translated from German version that used “ja, eher ja, eher nein, nein”). In the fMRI sample, subjects had to wait 4s (jittered) before the response options were shown. Direction of options (from left to right or vice versa) was random. Directly after decision, the ITI started. If subjects failed to make a decision within 4s, ITI started and trial was counted as missing. ca.: circa, RT: reaction time

**Table 1:** Sample characteristics, means and p-values calculated by two-sided permutation test.

| variable | HC group | se | GD group | se | p perm test |
| --- | --- | --- | --- | --- | --- |
| years in school | 10.87 | 0.22 | 10.77 | 0.22 | 0.837 |
| vocational school | 2.47 | 0.24 | 2.77 | 0.26 | 0.464 |
| net personal income | 1207.37 | 118.12 | 1419.67 | 174.51 | 0.272 |
| personal debt | 7166.67 | 2277.95 | 36166.67 | 11242.95 | <0.001 |
| Fagerström | 1.53 | 0.41 | 2.77 | 0.55 | 0.081 |
| age | 39.30 | 1.89 | 41.40 | 2.33 | 0.477 |
| AUDIT | 4.77 | 0.86 | 5.30 | 1.17 | 0.755 |
| BDI-II | 5.94 | 0.95 | 12.83 | 1.88 | 0.003 |
| SOGS | 1.87 | 0.54 | 9.17 | 0.57 | <0.001 |
| KFG | 3.70 | 1.05 | 28.47 | 1.54 | <0.001 |
| BIS-15 | 32.40 | 1.15 | 33.60 | 1.10 | 0.468 |
| GBQ persistence | 2.18 | 0.21 | 3.24 | 0.20 | 0.001 |
| GBQ illusions | 3.18 | 0.26 | 3.52 | 0.22 | 0.334 |
| ratio female | 0.30 | - | 0.23 | - | 1.000* |
| ratio unemployed | 0.10 | - | 0.30 | - | 0.217* |
| ratio smokers | 0.53 | - | 0.67 | - | 0.299* |
| ratio right-handed | 0.93 | - | 0.93 | - | 1.000* |

\*chi-square test used; se: bootstrapped standard errors; years in school: years in primary and secondary school; vocational school: vocational school and/or university; Fagerström: smoking severity. AUDIT: alcohol use severity; BDI II: depressive symptoms, SOGS: South Oaks Gambling Screen to check for pathological gambling; KFG: Kurzfragebogen zum Glücksspielverhalten, Short Questionnaire Pathological Gambling, German diagnostic tool and severity measure based on the DSM-IV; BIS: Barratt Impulsiveness Scale for impulsivity; GBQ persistence and GBQ illusions: from the Gamblers' Beliefs Questionnaire, collecting gambling related cognitive distortions (Supplements 1.1)

### Subjective cue ratings

After the task, subjects rated all cues using the Self-Assessment Manikin (SAM) assessment^36^ (reporting on valence: happy vs. unhappy, arousal: energized vs. sleepy, dominance: in control vs. being controlled) and additional visual analogue scales: 1) “How strongly does this image trigger craving for gambling?” 2) “How appropriately does this image represent one or more gambling games?” 3) “How appropriately does this image represent possible negative effects of gambling?” 4) “How appropriately does this image represent possible positive effects of gambling abstinence?”. All scales were operated via a slider from 0 to 100.

All cue ratings were z-standardized within subject. Ratings were analyzed one-by-one using linear mixed-effects regression, using lmer from the lme4 package in R^37^, where cue category (and clinical group) denoted the fixed effects and subjects and cues denoted the sources of random effects.

### Estimating subject-specific parameters from behavioral choice data

We modeled each subject’s behavioral data by submitting dichotomized choices (somewhat no, no: 0; somewhat yes, yes: 1) into logistic regressions. We dichotomized choices to increase the precision when estimating behavioral parameters, in line with previous studies using the mixed gambles task^23,35^. Regressors for subject-wise logistic regressions were gain (mean-centered) and absolute loss (mean-centered) from the mixed gamble, as well as gamble simplicity (*ed*), loss-gain ratio and cue category of the stimulus in the background of the mixed gamble. We defined different logistic regressions by using different trial-based definitions of gamble value (*Q*) (see **Tab. S1**), submitted to the logistic function:

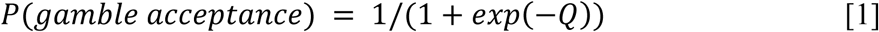

Different trial-based definitions of gamble value (*Q*) reflected two things:

1. Different ways of modeling LA may be adequate to distinguish a GD from a HC subject^23,25,27,35^ (**Tab. S1**).
2. Different ways of incorporating cue effects on decision-making (PIT effects) may be adequate to distinguish a GD from a HC subject. For example, the model **lac** assumes …

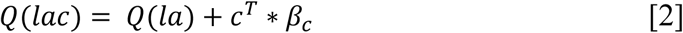

…where …

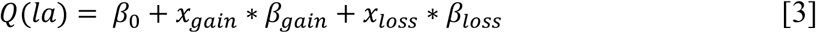

where *β*_*0*_ is the intercept, *x*_*gain*_ the objective gain value of the gamble, *β*_*gain*_ the regression weight for *x*_*gain*_ (same holds for *x*_*loss*_ and *β*_*loss*_, respectively), and c the dummy-coded column vector indicating the category of the current cue and *β c* a column vector holding the regression weights for the categories. **Lac** thus is a weighted linear combination of objective gain, objective loss with an additive influence of cue category. That is, some influence of cue category on decision-making (PIT) is modeled. Note that we have multiple PIT effects here, because *βc* is a vector of length three, reflecting the three affective categories (gambling, negative, positive) different from neutral. There were also models that did not incorporate any influence of loss aversion or category (intercept-only, **a**), or just of category (**ac**), or just of loss aversion (**la**) or of their interaction (**laci**):

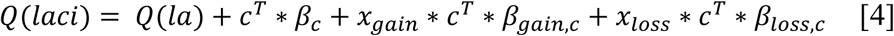

A model selection procedure could thus choose whether cue-induced effects on loss aversion (“loss aversion PIT”, i.e. the **laci** model) were important or not to distinguish between GD and HC subjects. Logistic regressions were fit using maximum likelihood estimation within the glm function in R^38^. Resulting regression parameters were extracted per model (e.g. *β*_*0*_, *β*_*gain*_, *β*_*loss*_ for model **la**) and subject. We appended the loss aversion parameter (*λ*) to the estimated coefficients by computing for each subject and pair of *β*_*gain*_, *β*_*loss*_:

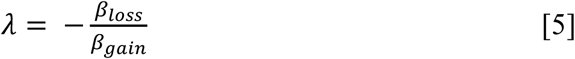

Models with names incorporating a “c” (e.g. **lac** or **laci**) are those that assume some influence of the cues (i.e. PIT effects). Models **laCh**, **laChci** are from^27^. Note that per model each subject thus had a characteristic *parameter vector* (the estimated regression weights, plus, in the expanded case, the loss aversion coefficients) and all subjects’ parameter vectors belonging to a certain model constituted the model’s *parameter set*. There were 13 different ways (i.e. models) to extract the behavioral parameters per subject plus 8 expansions by computing the loss aversion parameters after model estimation (**Tab. S1**), i.e. 21 parameter sets. In a separate analysis, the models were estimated with adjustment for cue repetition (using one additional two-level factor in each single-subject model) and by randomly selecting 45 gambling cues out of 67, to equalize the number of trials per cue category.

### Classification

Our machine learning approach is based on regularized regression and cross-validation as used in other machine learning studies in addiction and psychological research^30,31,39^.

#### Overall reasoning in building the classifier

The main interest of our study was to assess whether cue-induced changes in decision-making during an affective mixed gambles task can be used to distinguish GD from HC subjects. We hypothesized that shifts in loss aversion that depend on what cues are shown in the background (“loss aversion PIT”) should best distinguish between GD and HC subjects. This means, the **laci** model’s parameter set should have been the most effective in distinguishing between GD and HC subjects. To test this hypothesis, we used a machine learning algorithm based on regularized logistic regression that selected among various competing parameter sets (from the 21 different models, **la**, **lac**, **laci**, etc.) the set that best distinguished HC and GD subjects.

To assess the generalizability of the resultant classifier, we used cross-validation (CV)^30,32,39,40^. Generalizability estimates the predictive power, and hence replicability, of a classifier in new samples^28^. Note that machine learning algorithms are designed to generalize well to new samples by inherently avoiding overfitting to the training data^41(p9)^. We computed a p-value of the algorithm denoting the probability that its classification performance was achieved under a baseline model (predicting using only smoking severity as predictor variable).

Beyond cross-validation, which uses only one data set (splitting it repeatedly into training and test data set), validation of a classifier on a completely independent sample is the gold-standard in machine learning to assess the quality of an estimated model^28^. Hence, we estimated the generalization performance also via application of our classifier to a completely independent sample of HC and GD subjects, who had performed a slightly adapted version of the task in an fMRI scanner. A p-value was computed, as above, with random classification as the baseline model. For detailed information on estimating the classifier, please see **Supplements (1.4 and Fig. S1)**. For classical analyses of group comparisons regarding gamble acceptance rate and loss aversion parameters, please see **Supplements (1.6).** In a separate analysis, we ran the classification with the model parameters adjusted for cue repetition and with equalized number of trials per cue category.

### Ethics

Subjects gave written informed consent. The study was conducted in accordance with the Declaration of Helsinki and approved by the ethics committee of Charité – Universitätsmedizin Berlin.

## RESULTS

### Cue ratings

Gambling cues were seen as more appropriately representing one or more gambling games than any other cue category: gambling > neutral (β = 1.589, p < 0.001), gambling > negative (β = 1.197, p < 0.001), gambling > positive (β = 1.472, p < 0.001). They elicited more craving in GD subjects (β = 0.71, p < 0.001). Negative cues were seen as evoking more negative feelings in both groups (β = -0.775, p < 0.001) and were seen as representing negative effects of gambling, more than any other category (**Supplements 2.1**). Positive cues were indeed seen as more representative for positive effects of gamble abstinence than any other category (**Fig. S2**).

### Prediction of group using behavioral data

The classification algorithm yielded an AUC-ROC of 68.9% (under 0-hypothesis, i.e. with only smoking as predictor: 55.1%, p = 0.002) (**Fig. 2B, S4**). The most often selected model was the “acceptance rate per category” (**ac**) model (90.7% of the rounds). Combined with the models **laec, lac** in 95.8% of the rounds a model was used that incorporated PIT, i.e. an effect of cue category on decisions (**Fig. S5**). In only 9.3% of the rounds a model was selected that incorporated loss aversion (i.e. gain and loss sensitivities). Validating the estimated classifier in the independent sample, the classifier yielded an AUC-ROC of 65.0% (under random classification: 55.3%, p = 0.047) (**Fig. 2C**). Adjusting for cue repetition and equalizing the number of trials across cue categories lead to very similar AUR-ROC scores, the **ac** model was still the most often chosen model (42%), otherwise **laec_LA** and **lac** were chosen very often (**Supplements 2.4**).

**Figure 2:**
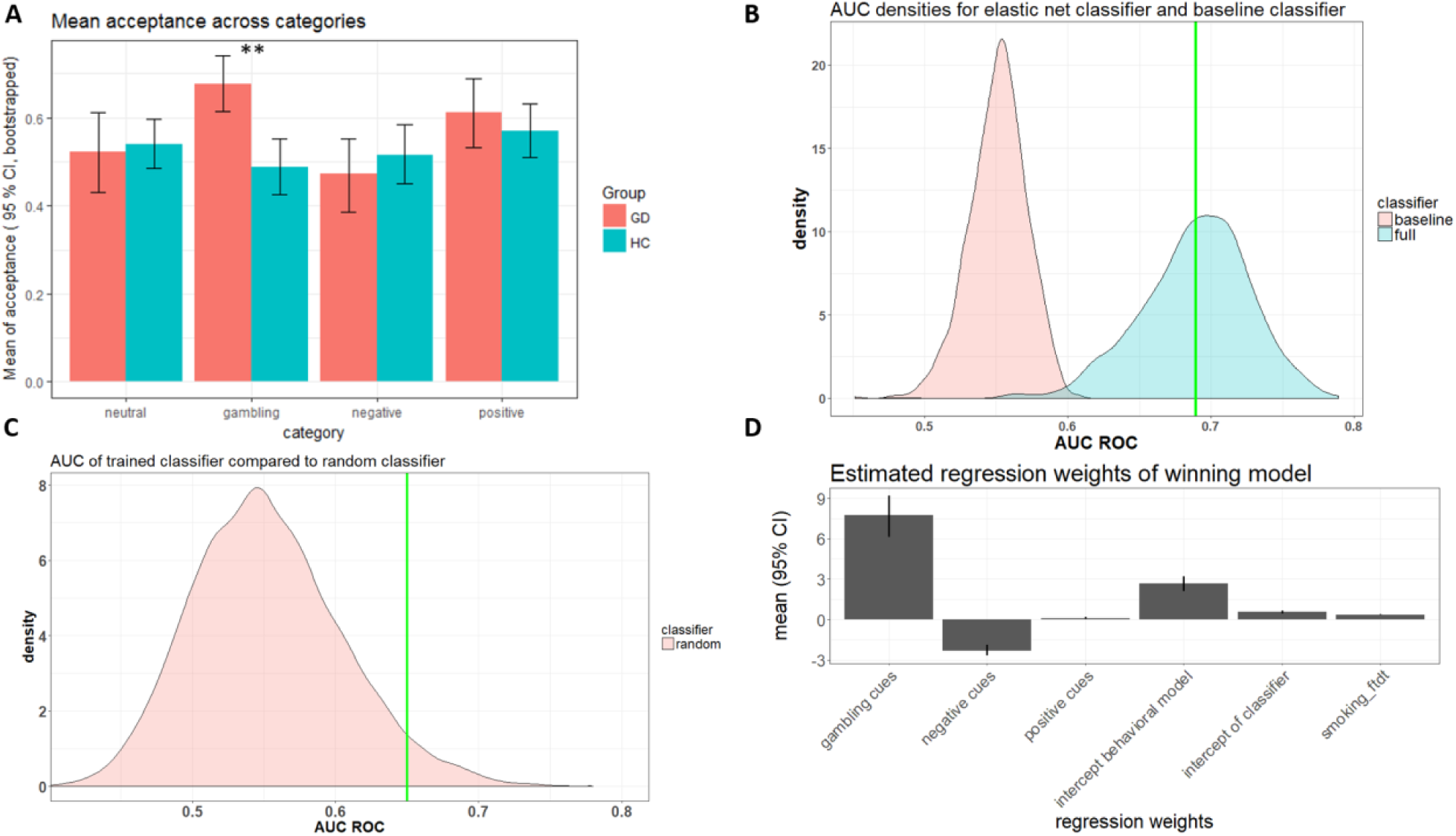
Behavioral results. A: Empirical mean acceptance rate with 95% CI’s. Means were computed over subjects’ means in the categories. Mean acceptance rate was significantly higher in GD subjects during gambling stimuli (p = 0.004). CIs are bootstrapped from category means of subjects. **B: Assessment of AUC-ROC of classifier:** Plot shows density estimates of the area under the receiver-operating curve when running the baseline classifier (red) / the full classifier (turquoise) 1000 times to predict the class label of 60 subjects. The green line shows the mean AUC performance of the estimated classifier across CV rounds. **C: Classifier validation on fMRI sample.** Plot shows the estimated density of AUC-ROC under random classification. The green line shows the performance of the combined 1000 classifiers on the fMRI data set. **D: Winning model for classification.** Standardized regression parameters and their confidence intervals (percentiles across cross-validation rounds). The algorithm mainly used the shift in acceptance rate in response to gambling cues in order to detect GD subjects.

### Inspection of classifier

Inspecting the classifier’s logistic regression weights, we saw that the classifier places most importance on the shift in gambling acceptance during gambling cues (see **Fig. 2D**). Note further that the classifier places also some importance on the sensitivity to the negative cues but deselects the sensitivity to positive cues.

### Acceptance rate and loss aversion under cue conditions

Overall acceptance rate between groups was not significantly different (HC: 53%, GD: 58%, p = 0.169, ΔAIC = 0). Across all subjects there was a significant effect of cue category on acceptance rate (p < 0.001, ΔAIC = 648), driven by the effect of positive and negative cues. There was a significant interaction with group (p = 0.002, ΔAIC = 9). There, GD subjects showed significantly higher acceptance rate during gambling cues than HC subjects (HC: 49%, GD: 68%, p_WaldApprox_ = 0.003) (**Fig. 2A**), and there were no more cue effects in the HC group and no other significant cue effect differences between HC and GD.

The fixed effects for gain sensitivity, absolute loss sensitivity, and LA over all trials for HC (0.26, 0.42, and 1.64) were descriptively larger than for GD (0.19, 0.22, and 1.13). Testing the interaction between group, gain, and loss (i.e. testing for difference of LA between groups) via nested model comparison, yielded p < 0.001, ΔAIC = 93, with sensitivity to loss being significantly smaller in GD subjects p_WaldApprox_ = 0.011. Loss aversion was significantly smaller in GD than in HC (p_perm_ < 0.001). Loss aversion shifts due to category did not differ between groups (**Supplements 2.2**).

## DISCUSSION

Gambling disorder (GD) is characterized by impaired decision making^4^ and craving in response to gambling associated images^9^. However, it is unclear whether specific cue-induced changes in loss aversion exist that distinguish GD from HC subjects. In order to better understand the basic mechanisms of impaired decision-making in addiction, we thus used a machine-learning algorithm to determine the relevance of cue-induced changes on loss aversion (“loss aversion PIT”) in distinguishing GD from HC subjects. We hypothesized that cue-induced changes in gamble acceptance and especially a strong shift of loss aversion by gambling and other affective cues should distinguish GD from HC subjects (i.e. the model representing this effect should have been chosen most often by the algorithm to distinguish GD from HC subjects). To our knowledge, our study is the first to investigate the classificatory power of addiction-relevant behavioral task parameters when distinguishing GD from HC subjects. Moreover, we are not aware of any study specifically investigating the relevance of behavioral PIT effects in characterizing addicted subjects using predictive modeling.

Our algorithm was significantly better in distinguishing GD from HC subjects than the control model, which only used smoking severity as a predictor variable (cross-validated AUC-ROC of 68.9% vs. 55.1%, p = 0.002). In an independent validation sample the classifier was almost as accurate (AUC-ROC of 65.0% vs. 55.3%, p = 0.047). When classifying subjects, in 93% of the estimation rounds, our algorithm chose a model with some influence of the cue categories on choices. The most frequently chosen model was the **ac** model (85%), i.e. a model only accounting for mean shifts in acceptance rate depending on cue category. PIT-related variables could therefore successfully discriminate between GD and HC subjects. We saw that especially the tendency of subjects to gamble more during the presentation of gambling cues was indicative of the subject belonging to the GD group. Contrary to what we expected, “loss aversion PIT” was not useful in distinguishing between GD and HC subjects. In other words, the algorithm never selected the **laci** model, which included the modulation of gain and loss sensitivity by cue categories. We also did not see this in univariate group comparisons. “Loss aversion PIT” might thus not play a role in distinguishing GD from HC subjects. However, small sample size, as in the present study, may limit the possible complexity of a classifier^42(p237)^. It cannot be ruled out that larger and more diverse samples in future studies may produce classifiers allocating at least minor importance to “loss aversion PIT”.

We observed that both GD and HC subjects perceived the cues as intended. GD subjects reported higher craving for gambling in response to gambling stimuli as seen in other studies^9^. Our results may thus be interpreted as cue reactivity leading to more automatic decision-making in GD subjects. Note that this does not mean that GD subjects simply show higher vigor or more disinhibition to press a button, as in some PIT designs^43^. Instead, since the required motor response for saying yes or no changed randomly, gamblers seemed to be indeed more inclined to *decide* in favor of gambling when gambling cues were shown in the background. Especially because cue influence on LA was not relevant for distinguishing GD from HC subjects, but instead cue influence on general acceptance rate, this may be seen as GD subjects responding more habitually and in a less goal-directed manner^15^ when gambling cues are visible.

In the current study, the classifier also put some importance on behavior under negative cues, and, descriptively but not significantly, GD subjects tended to reduce gambling more in the face of negative cues than HC subjects. Future studies should explore the possible power of negative images to inhibit gambling in larger and more heterogeneous GD samples.

Our results show the gambling promoting effects of gambling cues in GD subjects. Alcohol and tobacco advertisement promote alcohol and tobacco use^44^ and advertisement bans and counter-active labels on alcohol and tobacco goods help reduce consumption^45^. Our results suggest that much like advertisement for these substances, visual stimuli in gambling halls and on slot machines may also increase PIT effects. Policy makers may consider our results as another piece of evidence that gambling advertisement is not different from alcohol and tobacco advertisement and that respective advertisement regulation perhaps should be extended.

We are not aware of any machine learning studies that have assessed the relevance of a behavioral task measure in characterizing GD. Using this approach, we observed a cross-validated classification performance of AUC-ROC = 0.68. We are aware of one machine learning study that built and tested a classifier in 160 GD patients and matched controls based on personality questionnaire self-report, reaching an AUC-ROC = 0.77^31^. Studies in the field of substance-based addiction, using behavioral markers and machine learning for classification, report cross-validated AUC-ROC’s of 0.71 to 0.90 for cross-validated classification performance^30,39^. However, the mentioned studies used a whole array of different informative variables while the current studied tried to carve out the relevance of one basic behavioral mechanism while controlling for all covariates of no-interest.

Our results may be a first building block in creating more advanced and more multivariate diagnostic tools for GD and other addictive disorders, especially when combined with other high-performing discriminating features, such as personality profiles and scores from other decision-making tasks. Further, our results invite more in-depth scrutiny of decision-making in GD subjects during the presence of cues, e.g. on neural level^34^. Moreover, the above machine learning studies did not use an independent validation sample to corroborate their results. Our independent validation yielded an AUC-ROC of 0.65. This supports the validity of our findings of increased PIT in GD.

## STRENGTHS AND LIMITATIONS

When carving out the relevance of PIT, we did not match for depression score (BDI) because, epidemiologically, GD is associated with high depression scores^46^, meaning it could be seen as a feature of GD. Further, the evidence on the association of PIT and depression is inconclusive^47,48^. However, PIT might play some role in depression and thus also in GD subjects. Future studies should thus address the modulatory effect of depressive symptoms in GD on PIT^49^.

The current classifier was slightly less effective in the independent validation sample than estimated using cross-validation (AUC = 65.4% vs. 68.0%). This might have occurred due to the use of an fMRI version of the affective mixed gambles task in the validation sample. It included an additional decision-making period, during which subjects could not yet answer. This may have led to slight changes in responses with respect to the cue categories. However, this could be seen as a strength since our classifier still performed better than chance. And classifiers that are robust against slight changes in the experimental set-up allow arguably more general conclusions than classifiers that only work with data from the same experimental set-up. Future studies should also use validation samples^40^.

Cues were repeated and trial numbers were not perfectly balanced across categories. We adjusted for this in our analyses and results were stable. Here, model selection geared also towards reduced loss aversion additionally characterizing GD, in line with^23,24^.

## CONCLUSION

Our results propose that GD subjects’ acceptance of mixed gambles is cue-dependent and that this cue-dependency even lends itself to distinguishing GD from HC subjects in out-of-sample data. However, we did not observe that cues specifically shift loss aversion, neither on average, nor in a way relevant to classification. We saw that especially gambling cues lead to increased gambling GD subjects. Observing increased PIT in GD suggests that PIT related to an addictive disorder might not depend on the direct effect of a substance of abuse, but on related learning processes^17^ or on innate traits^18^. The here reported effects should be explored further in larger, more diverse and longitudinal GD samples as they could inform diagnostics, therapy^50^ and public health policy.

## ONLINE MATERIAL

You can find the data and R Code to reproduce the analyses here: https://github.com/pransito/PIT_GD_bv_release

## AUTHORS’ CONTRIBUTION

AG designed the experiment, collected the data, analyzed the data, and wrote the manuscript. MA implemented the ratings and questionnaire electronically, analyzed the ratings data, and revised the manuscript. KB collected data and revised the manuscript. CM reviewed the machine-learning algorithm and revised the manuscript. AH revised the manuscript, and oversaw manuscript drafting and data analyses. AW revised the manuscript and oversaw implementation of experiment in the lab. NK revised the manuscript and, advised first author. NRS designed and supervised study and experiment, and oversaw manuscript drafting and data analyses.

