## Supplemental Materials for "Cue-induced effects on decision-making distinguish subjects with gambling disorder from healthy controls"

**Title of article:**

### 1 Supplementary methods

#### 1.1 Sample

We recruited GD subjects via eBay classifieds, and notices in Berlin casinos and gambling halls. Any known history of a neurological disorder or a current psychological disorder (except tobacco dependence) as assessed by the Screening of the Structured Clinical Interview for DSM-IV Axis I Disorders (SCID-I) (First et al., 2002) lead to exclusion from the study. There were five subject dropouts (two technical error, one rejected all gambles, two to improve matching). The final sample consisted of 30 GD and 30 HC subjects (**Tab. 1**). According to the South Oaks Gambling Screen (Lesieur and Blume, 1987; Stinchfield, 2002) (3-point Likert scales), GD subjects differed in gambling habits to HC mainly in frequency of playing slot machines (most frequent answer of GD: “3: once a week or more”, HC: “1: not at all”) ($t = 7.30$, $p < 0.001$) and casinos (most frequent answer of GD: “3: once a week or more”, HC: “1: not at all”) ($t = 3.99$, $p = 0.001$). 21 GD indicated “3: once a week or more” for slot machines, 12 indicated that answer for casinos and 6 for sports betting. Further these questionnaires were used: Fagerström: smoking severity (Heatherton et al., 1991); AUDIT: alcohol use disorders identification test to test alcohol use severity (Dybek et al., 2006); BDI II: Beck’s Depression Inventory to check for depressive symptoms (Beck et al., 1996), KFG: Kurzfragebogen zum Glücksspielverhalten, Short Questionnaire Pathological Gambling, German diagnostic tool and severity measure based on the DSM-IV (Petry and Baulig, 1996); BIS: Barratt Impulsiveness Scale for impulsivity (Meule et al., 2011); GBQ persistence and GBQ illusions: from the Gamblers’ Beliefs Questionnaire, collecting gambling related cognitive distortions (Steenbergh et al., 2002)

#### 1.2 Cue selection

For the purposes of this study four sets of images were assembled: 1) 67 gambling images, showing a variety of gambling scenes, situations and cues: 36 showing different kinds of slot machines, 12 showing poker, 13 showing roulette, 3 featuring money, 3 featuring dice; 2) 31 images showing negative consequences of gambling (as of now referred to as *negative images*): 7 showing depression / sadness, 4 depicting poverty, 4 depicting debt, 3 showing a quarrel between people, 2 showing family problems, 2 showing the lack of money, 2 symbolizing suicide, 2 showing money burning; 3) 31 images showing positive effects of abstinence from gambling (as of now referred to as *positive images*): 6 showing family, 4 showing relationships, 4 showing friendships, 3 depicting success, 3 depicting freedom, 3 showing joy, 2 showing saved money; 4) 24 neutral images showing objects: 6 kitchen utensils, 8 showing other household objects, 2 showing tools, 2 showing abstract paintings. None of the neutral pictures showed humans or faces.

Images were obtained from the internet, sought purposefully to fit the defined categories (positive, gambling, neutral). Online search for images was performed using popular image search engines. Groups of selected images were matched for content as follows: a) percent of images showing a social stimulus (i.e. a person) as opposed to images without persons (gam: 88.2%, pos: 77.4%, neg: 90.3%; $Chi^2 = 1.123$, df = 2, p = 0.570); b) percent of images showing a face as opposed to people with their face turned away or just hands (gam: 35.3%, pos: 38.7%, neg: 51.6%; $Chi^2 = 3.530$, df = 2, p = 0.171); c) percent of images showing males (gam: 67.6%, pos: 64.5%, neg: 77.4%; $Chi^2 = 1.300$, df = 2, p = 0.523).

All images were cropped to fit the aspect ratio optimized to minimize the loss of image area (3:2). Each image was cropped individually making sure that no content was lost. All the images were resized to the resolution of the lowest image in the set (450 x 300 pixels), ensuring that the image dimensions and quality are the same across all images. The images can be acquired with the corresponding author for scientific purposes upon reasonable request. Due to copyright issues they cannot be shared publicly.

#### 1.3 Behavioral models

**Table S1: Definition of gambling value.**

| name | definition of value in each trial | np | nep |
|---|---|---|---|
| **a** | $Q(a) = \beta_0$ | 1 | - |
| **lar** | $Q(lar) = \beta_0 + x_{ratio} \cdot \beta_{ratio}$ | 2 | - |
| **laCh** | $Q(laCh) = x_{gain} \cdot \beta_{gain} + x_{loss} \cdot \beta_{loss}$ | 2 | 3 |
| **la** | $Q(la) = \beta_0 + x_{gain} \cdot \beta_{gain} + x_{loss} \cdot \beta_{loss}$ | 3 | 4 |
| **ac** | $Q(ac) = c^T \cdot \beta_c$ | 4 | - |
| **lae** | $Q(lae) = Q(la) + ed \cdot \beta_{ed}$ | 4 | 5 |
| **larc** | $Q(larc) = Q(lar) + c^T \cdot \beta_c$ | 5 | - |
| **lac** | $Q(lac) = Q(la) + c^T \cdot \beta_c$ | 6 | 7 |
| **laec** | $Q(laec) = Q(la) + ed \cdot \beta_{ed} + c^T \cdot \beta_c$ | 7 | 8 |
| **larci** | $Q(larci) = Q(larc) + x_{ratio} \cdot c^T \cdot \beta_{ratio,c}$ | 8 | - |
| **laChci** | $Q(laChci) = Q(laCh) + x_{gain} \cdot c^T \cdot \beta_{gain,c} + x_{loss} \cdot c^T \cdot \beta_{loss,c}$ | 8 | 12 |
| **laci** | $Q(laci) = Q(la) + c^T \cdot \beta_c + x_{gain} \cdot c^T \cdot \beta_{gain,c} + x_{loss} \cdot c^T \cdot \beta_{loss,c}$ | 12 | 16 |
| **laeci** | $Q(laeci) = Q(la) + ed \cdot \beta_{ed} + c^T \cdot \beta_c + x_{gain} \cdot c^T \cdot \beta_{gain,c} + x_{loss} \cdot c^T \cdot \beta_{loss,c}$ | 16 | 20 |

$x_{sub}$: independent variable; $\beta_{sub}$: regression weight, i.e. free parameter of model c: dummy coded cue category variable as vector; T: transpose; np: number of parameters in model; nep: number of parameters if model parameters were expanded by post-hoc computation of loss aversion parameters; the collection of parameter vectors for all subjects for one model is the parameter set of that model; adding to the 13 “np” parameter sets the 8 “nep” parameter sets, we thus have 21 parameter sets; a: model with intercept only, i.e. mean acceptance over all trials, ac: acceptance rate per category; ed, Gamble simplicity is defined as Euclidean distance from diagonal of gamble matrix (ed) (Tom et al., 2007)

**Lar*** models are ratio models where only predictor for gamble value is *ratio = gain/loss* for each trial (Gelskov et al., 2016). **La** model is the classical loss aversion model (Genauck et al., 2017; Tom et al., 2007). **Lae** is the model when adding *ed* (gamble simplicity) as additional predictor model (Genauck et al., 2017; Tom et al., 2007). **Lac** adds the category as additional linear effect (modulation of intercept). **Laci** adds further an interaction of gain and loss sensitivity (and thus with loss aversion) with category. **Laec** and **Laeci** are the same expect they add gamble simplicity again. The model **a** only estimates the intercept and **ac** only the shift of intercept depending on cue category (gambles are ignored). **laCh** is the De Martino/Charpentier model (Charpentier et al., 2016; De Martino et al., 2010), $Q(laCh) = 1*gain + \lambda*loss$), subjected to a two-options softmax function $P(accept = 1) = (1+\exp(-\mu*value))^{-1}$

with $\mu$ as a free parameter (i.e. the general case of a logistic function). Note, however, that **laCh**'s value function can be rewritten as $Q(laCh) = \mu * gain + \mu * \lambda * loss$, which then is submitted to the logistic function (i.e. two-option softmax function without any free parameter). **laCh** is hence a logistic regression like **la** but without an intercept $\beta_0$, with $\beta_{gain} = \mu$ and $\beta_{loss} = \mu * \lambda$ (hence $\lambda = \beta_{loss} / \beta_{gain}$). Hence, **laChci** (Charpentier et al., 2016) can be formed accordingly.

#### 1.4 Classification using behavioral data

##### *1.4.1 Detailed description of algorithm to build classifier*

From 21 different parameter sets (**Tab. S1**), representing different "loss aversion PIT" (e.g. the **laci** model) and respective control models, we wanted to find the best parameter set to build a classifier (here a logistic regression model) to distinguish between GD and HC subjects in out-of-sample test data (Guggenmos et al., 2018; Whelan et al., 2014) (**Fig. S3**). We expected to see the **laci** model winning, because it assumes an interaction between loss aversion and cue categories.

In a first step we used model selection based on cross-validation to find the parameter set that best distinguished between GD and HC (Arlot and Celisse, 2010; Bratu et al., 2008; Varma and Simon, 2006). Using cross-validation for model selection ensures that the selected model will be the one that best generalizes to out-of-sample data (and hence overfitting is avoided). The algorithm used the different parameter sets, one by one, to predict group membership of subjects using logistic ridge regression (Le Cessie and Van Houwelingen, 1992) **(Section 1.5)**. Ridge regression has one hyperparameter that is tuned to optimize the cross-validated classification power of each parameter set according to the area under the receiver operating curve (AUC-ROC) (Ahn et al., 2016; Ahn and Vassileva, 2016; Whelan et al., 2014; Zacharaki et al., 2009).

AUC-ROC ranges from 0.5 (chance) to 1 (perfect sensitivity and specificity) (Provost et al., 1998). The parameter set with the highest cross-validated AUC-ROC was selected.

In a second step, the classifier was completed. Smoking severity was not entirely matched between the groups. This is why the algorithm added smoking severity to the parameter set with the best cross-validated AUC-ROC score from the first step to logistic elastic net regression (Zou and Hastie, 2005) **(Section 1.5)**, optimizing the AUC-ROC by tuning its two hyper-parameters (Whelan et al., 2014), again via cross-validation (**Fig. S3**). We did not use elastic net regression during model selection because it can force regression parameters to zero (sparse models) (Zou and Hastie, 2005). This would have blurred the interpretable differences between the behavioral models. However, we used elastic net regression in the last step of classifier building, because we were interested whether the algorithm would force parameters of the winning model from the first model selection step to zero, e.g. because a parameter does not add any more information to classification.

We assessed the generalizability of the above algorithm 1000 times via 10-fold cross-validation (Arlot and Celisse, 2010), which yielded a distribution of classifiers and thus of AUC-ROC's. Note that the cross-validation to estimate generalizability lead to the cross-validations used in the machine learning (for tuning of hyperparameters) of the first and second step to become *nested*, which is necessary to avoid contamination between training and test data (Arlot and Celisse, 2010; Bratu et al., 2008; Varma and Simon, 2006; Whelan et al., 2014). We computed the mean of the obtained AUC-ROC's and estimated its p-value by performing the exact same 1000 CV rounds but each time with only smoking severity as predictor (baseline classifier). We then subtracted the AUC-ROC's of the baseline classifiers one-by-one from the 1000 AUC-ROC's of the full classifiers. This yielded a distribution of classification improvement (i.e., improvement of AUC-ROC due to using the full classifier instead of the baseline classifier).

We tested this distribution against the value of classification improvement under the null-hypothesis (i.e. zero improvement) to obtain a p-value of significance of classification improvement.

To build the final interpretable and reportable classifier, one would usually apply the algorithm *once* to the complete data set. Since the application to the complete data set still entails cross validation for tuning of the ridge and elastic net regressions' hyperparameters leading to slightly varying classifiers, the algorithm was not run once but 1000 times on the complete data set. We plotted the ensuing distribution of selected parameter sets and the distribution of the respective regression weights as per-parameter means with 95% percentile bounds. For a graphical illustration of the algorithm see **Fig. S3**. For the R code and the data please see https://github.com/pransito/PIT_GD_bv_release.

##### *1.4.2 Validating the classifier on an independent sample*

We applied all 1000 classifiers estimated on the full data set to each of the 60 subjects of the validation sample, yielding 1000 decision values per subject (real-valued scalars). To incorporate the complete distribution of the classifiers, we summed up, for each subject, the decision responses of all 1000 estimated classifiers, yielding one decision value per subject. Using the known true labels of all subjects, the decision values of all subjects were used to compute the AUC-ROC. We compared this obtained AUC-ROC to its distribution under a null-model (10,000 repetitions of random, i.e. coin-flip, classification), to compute a p-value.

### PREDICTION OF GROUP

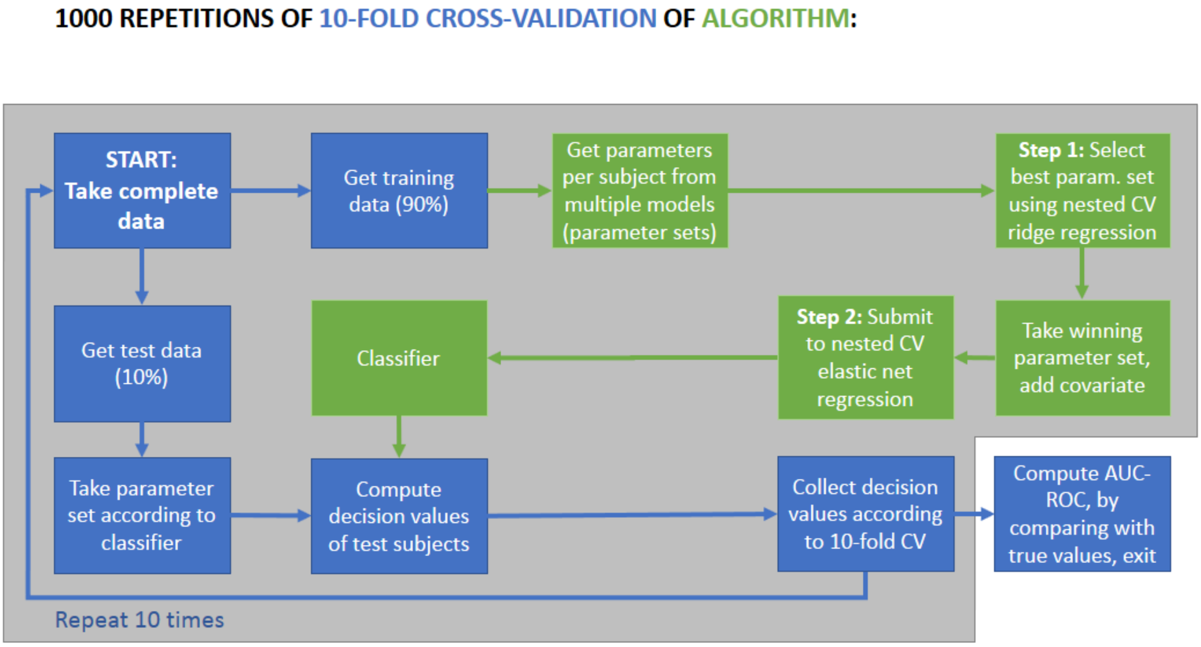


**Figure S1: Classification algorithm (in green) and its cross-validation (in blue).** 10-fold-cross-validation of the algorithm that builds a classifier to predict the group (i.e. label) of each subject (healthy control vs. subjects with gambling disorder) based on behavioral data. On each fold, data was split into training and test set. Training had 90% (54), test had 10% (6) data points. As preparation, all subjects have their behavioral choice data modeled according to the 21 candidate models, leading to 21 parameter sets. **Step 1:** All parameter sets are one-by-one used to predict group membership using logistic ridge regression. Performance is assessed using nested 5-fold cross-validation (CV), i.e. training has 72% (43) data points and test has 18% (11) data points. Logistic ridge regression is performed over a range of values of the penalty parameter to get optimal nested CV performance per parameter set. The best performing parameter set is declared winner. If there are ties, the simpler parameter set is declared winner. For stability, this step is repeated 10 times and the most often winning model is forwarded. **Step 2:** The winning parameter set plus covariate “smoking severity” is subjected to a logistic elastic net regression optimizing for its two hyper-parameters using nested 10-fold CV (training: 81%, 49 data points; test: 9%, 5 data points). This gets repeated 10 times for stability and from 10 models a mean model is computed. This yields a classifier, i.e. here a logistic regression model, which is applied to the initial 10% test data points. The procedure is repeated until all data points have been test data once and decision-values could be collected for all 60 subjects. The area under curve (AUC) under the receiver-operating (ROC) curve is then computed as CV score of interest. To account for the multiple possibilities of slicing the data into training and test for 10-fold CV and to compute a p-value the whole procedure was repeated 1000 planned times: a) running all steps described above; b) running only step 2 with only smoking severity as predictor (baseline model). All data splits ensured balanced labels (50/50) in training and test sets.

#### 1.5 Logistic elastic net regression

Logistic elastic net regression expands normal logistic regression by penalizing complicated regression solutions (large regression weights). How much it penalizes is governed by two hyper-paramters: λ and α, introduced in its expanded error (or cost) function (i.e. the measurement of how far off the fitted model's predictions are from the real data's labels):

$$L = COST(h(x), y) + \lambda[(1-\alpha)|\beta|_2^2/2 + \alpha|\beta|_1]$$

...where COST is the cross-entropy function (i.e. in short the negative log-likelihood of the model, (Bishop, 2006, 205ff.) exact equation of which is not relevant here. Further, *h(x)* is the model yielding a decision-value/a prediction, and x is the vector of predictors.

The model *h(x)* is a regression equation that can be written as $\theta(\beta^T x)$, where $\beta^T x$ is the scalar product of the regression weights stored in vector $\beta$and the predictor vector $x$, and $\theta$is the logistic transfer function. The regularization term $+\lambda[\ldots]$adds the size of the vector beta to the cost because it is a measure of complexity. Elastic net regression uses two measures, the L1-norm ($|\beta|_1$) and the L2-norm ($|\beta|_2$) and mixes them depending on the two hyperparameters $\lambda$ and $\alpha$. Upon estimation of $\beta$, L is minimized given λ and α (Zou and Hastie, 2005). Which hyperparameters to choose is a matter of tuning, e.g. via nested cross-validation. Ridge regression is a special case of elastic net regression, namely when α = 0.

#### 1.6 Group comparisons regarding acceptance rate and loss aversion parameters

To provide a fuller overview of the data, we also performed classical mean-differences analyses to analyze the choice data. We explored the effect of the independent variables "cue category" and "group" onto the dichotomized dependent variable "choice". We thus fitted logistic linear mixed effects models using R's lme4 package (Bates et al., 2015) with "cue category" and "group" as sources of fixed effects and "subject" and "cue" as sources of random effects. Concerning LA, we explored the effect of the independent variables "gain" and "loss" onto the dependent variable "choice". We thus fitted a logistic linear mixed effects model with "gain" and "loss" as sources of fixed effects and "subject", "cue" and "cue category" as sources of random effects. We tested for significance of the effects of independent variables using nested-models chi-square-difference tests (i.e. likelihood-ratio tests) and ΔAIC (with positive ΔAIC meaning an improvement in model fit). We performed further model comparisons with models successively incorporating higher-order interactions of the independent variables (("gain" + "loss") X "cue category").

#### 1.7 Validation sample

**Table S2: Sample characteristics, means and p-values calculated by two-sided permutation test.**

| Variable | HC (N = 30) | se | PG (N = 30) | se | pooled se | p perm test |
|---|---|---|---|---|---|---|
| **years in school** | 10.87 | 0.19 | 10.13 | 0.24 | 0.21 | 0.031 |
| **vocational school** | 2.73 | 0.29 | 2.07 | 0.25 | 0.27 | 0.108 |
| **net personal income** | 1029 | 92.27 | 1106 | 138.93 | 115.6 | 0.667 |
| **personal debt** | 8500 | 3397 | 24000 | 9590 | 6494 | 0.097 |
| **Fagerström** | 1.97 | 0.43 | 3.03 | 0.51 | 0.47 | 0.138 |
| **age** | 35.37 | 1.66 | 37.37 | 2.01 | 1.84 | 0.459 |
| **AUDIT** | 4.8 | 0.59 | 4.87 | 1.05 | 0.82 | 1 |
| **BDI-II** | 5.1 | 1.03 | 11.57 | 1.72 | 1.38 | 0.002 |
| **SOGS** | 1.73 | 0.47 | 8.8 | 0.67 | 0.57 | < 0.001 |
| **KFG** | 2.37 | 0.74 | 35 | 1.64 | 1.19 | < 0.001 |
| **BIS-15** | 31.8 | 0.99 | 36.33 | 1.08 | 1.03 | 0.004 |
| **GBQ persistence** | 1.96 | 0.2 | 3.28 | 0.19 | 0.2 | < 0.001 |
| **GBQ illusions** | 2.41 | 0.24 | 3.73 | 0.22 | 0.23 | < 0.001 |
| **ratio female** | 0.20 | - | 0.20 | - | - | 1.000 |
| **ratio unemployed** | 0.17 | - | 0.20 | - | - | 1.000 |
| **ratio smokers** | 0.60 | - | 0.77 | - | - | 0.262 |
| **ratio right-handed** | 0.97 | - | 0.84 | - | - | 0.204 |

*chi-square test used; se: bootstrapped standard errors; years in school: years in primary and secondary school; vocational school: vocational school and/or university; Fagerström: smoking severity (Heatherton et al., 1991); AUDIT: alcohol use disorders identification test to test alcohol use severity (Dybek et al., 2006); BDI II: Beck's Depression Inventory to check for depressive symptoms (Beck et al., 1996), SOGS: South Oaks Gambling Screen to check for pathological gambling according to DSM-III (Lesieur and Blume, 1987); KFG: Kurzfragebogen zum Glücksspielverhalten, Short Questionnaire Pathological Gambling, German diagnostic tool and severity measure based on the DSM-IV (Petry and Baulig, 1996); BIS: Barratt Impulsiveness Scale for impulsivity (Meule et al., 2011); GBQ persistence and GBQ illusions: from the Gamblers' Beliefs Questionnaire, collecting gambling related cognitive distortions (Steenbergh et al., 2002)

### 2 Supplementary results

#### 2.1 Cue ratings

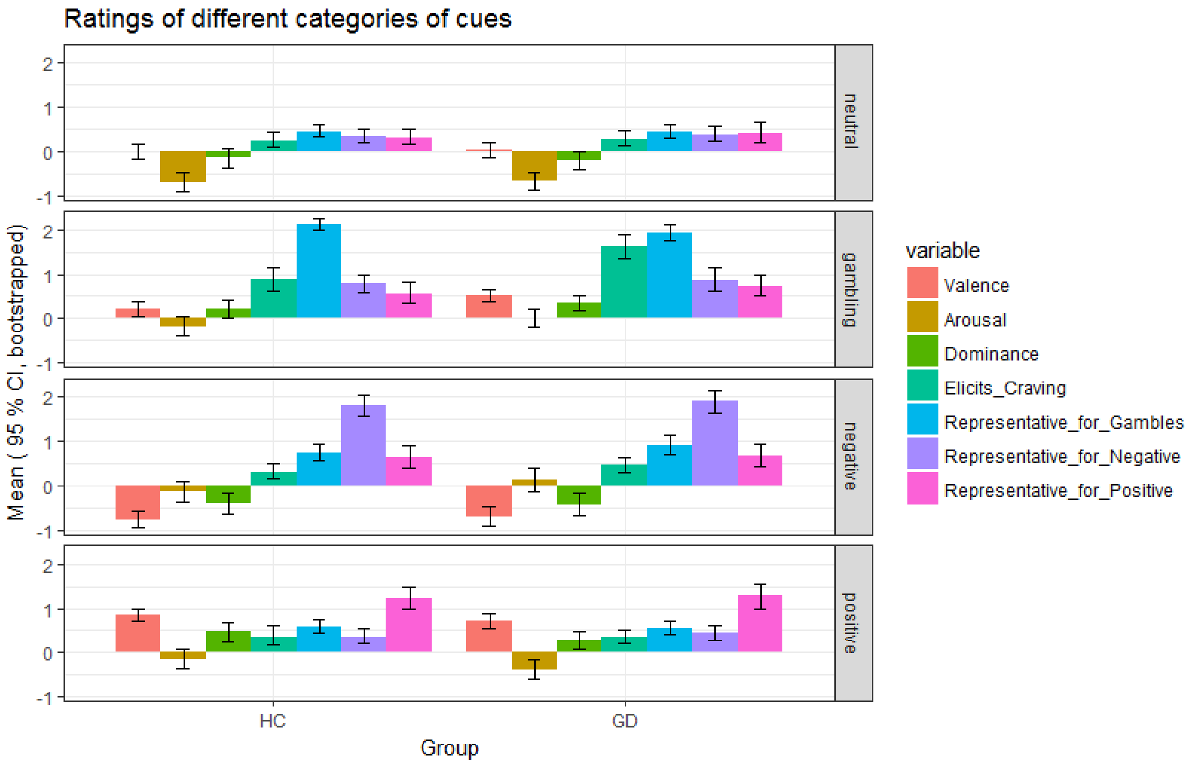


**Figure S2: Means and bootstrapped 95% confidence intervals of rating variables.** GD: subjects with gambling disorder, HC: healthy controls. Plot facets report from top to bottom on ratings of neutral category cues, gambling, negative and positive category cues. Neutral cues were indeed rated as neutral in valence and as eliciting low arousal (Lang et al., 1993)

##### 2.1.1 Valence

As expected, image category affected valence ratings ($\Delta Chi^2 = 1513$, $\Delta df = 11$, $p < 0.001$), where negative images were rated as lower than neutral images in valence ($\beta = -0.760$, $p < 0.001$), the positive category was rated higher in valence than neutral images ($\beta = 0.782$, $p < 0.001$).

Beyond image category group had a modulatory influence on valence ratings ($\Delta Chi^2 = 11$, $\Delta df = 4$, $p < 0.024$). PG subjects showed a trend in rating gambling pictures higher than HC subjects ($\beta = 0.268$, $p = 0.086$).

##### 2.1.2 Arousal

As expected, image category affected arousal ratings ($\Delta Chi^2 = 2067$, $\Delta df = 11$, $p < 0.001$), where neutral images were rated as lower than 0 in arousal ($\beta = -0.694$, $p < 0.001$). All other categories were rated as more arousing than neutral images (βs ranging from 0.428 to 0.698, all p's < 0.001). Gambling images were slightly more arousing than positive images ($\beta = 0.175$, $p = 0.023$). Negative images were slightly more arousing than positive images ($\beta = 0.270$, $p = 0.012$).

Beyond image category group had some modulatory influence on arousal ratings ($\Delta Chi^2 = 10$, $\Delta df = 4$, $p = 0.048$). PG subjects did not find gambling pictures more arousing than HC.

##### 2.1.3 Dominance

As expected, image category affected dominance ratings ($\Delta Chi^2 = 1963$, $\Delta df = 11$, $p < 0.001$), where the positive ($\beta = 0.552$, $p < 0.001$) and gambling ($\beta = 0.456$, $p < 0.001$) images were both rated higher than 0. Negative images ($\beta = -0.236$, $p = 0.013$) and neutral images ($\beta = -0.179$, $p = 0.023$) were rated as lower than 0 in dominance.

Beyond image category group did not have an influence on dominance ratings. ($\Delta Chi^2 = 4$, $\Delta df = 4$, $p = 0.352$)

##### 2.1.4 Craving inducing

As expected, image category affected craving ratings when both groups were combined ($\Delta Chi^2 = 3430$, $\Delta df = 11$, $p < 0.001$), respectively. Gambling pictures induced more craving for gambling than any other image category: gambling > neutral ($\beta = 0.999$ $p < 0.001$), gambling > negative ($\beta = 0.881$, $p < 0.001$), gambling > positive ($\beta = 0.901$, $p < 0.001$).

Beyond image category group had a significant influence on craving ratings. ($\Delta Chi^2 = 20$, $\Delta df = 4$, $p < 0.001$). PG subjects showed higher ratings for craving on gambling pictures ($\beta = 0.707$, $p < 0.001$) compared to HC subjects.

##### 2.1.5 Representativeness for gambling

As expected, image category affected gambling representativeness ratings ($\Delta Chi^2 = 684$, $\Delta df = 11$, $p < 0.001$). Gambling category images were more representative of gambling than any other category: gambling > neutral ($\beta = 1.606$, $p < 0.001$), gambling > negative ($\beta = 1.209$, $p < 0.001$), gambling > positive ($\beta = 1.482$, $p < 0.001$). Beyond image category group had no modulatory influence.

##### 2.1.6 Representativeness for negative effects of gambling

As expected, image category affected ratings of representativeness for negative effects of gambling ($\Delta Chi^2 = 1952$, $\Delta df = 11$, $p < 0.001$).

Negative image category was more representative of negative effects of gambling than any other group: negative > gambling ($\beta = 1.049$, $p < 0.001$), negative > positive ($\beta = 1.471$, $p < 0.001$), negative > neutral ($\beta = 1.514$, $p < 0.001$). Beyond image category group did not have an influence on ratings of representativeness for negative effects of gambling ($\Delta Chi^2 = 2$, $\Delta df = 4$, $p = 0.971$).

##### 2.1.7 Representativeness for positive effects of gambling abstinence

As expected, image category affected ratings of representativeness for positive effects of gambling abstinence ($\Delta Chi^2 = 2590$, $\Delta df = 11$, $p < 0.001$, both groups combined). The positive category was more representative of positive effects of abstinence from gambling than any other category: positive > neutral ($\beta = 0.889$, $p < 0.001$), positive > negative ($\beta = 0.590$, $p < 0.001$) and positive > gambling ($\beta = 0.605$, $p < 0.001$). Beyond image category group did not have an influence on ratings of representativeness for positive effects of gambling abstinence ($\Delta Chi^2 = 1$, $\Delta df = 4$, $p = 0.836$).

##### 2.1.8 How much do you question your gambling when seeing this image

This question was only answered by gamblers. As expected, image category affected the motivation of questioning the own gambling behavior ($\Delta Chi^2 = 1514$, $\Delta df = 11$, $p < 0.001$), where the negative images were rated higher than any other image category: negative > neutral ($\beta = 1.121$, $p < 0.001$), negative > positive ($\beta = 0.535$, $p < 0.001$) and negative images were also rated higher than gambling images ($\beta = 0.514$, $p = 0.003$). One subject did not answer these questions and hence was not included in this analysis.

#### 2.2 Group comparisons loss aversion models

Gain and loss had a significant influence on gamble choice in all subjects ($p < 0.001$, $\Delta AIC = 4414$). There was a significant fixed effect interaction with group that improved model fit ($p < 0.001$, $\Delta AIC = 93$). Gain, absolute loss sensitivity, and LA over all trials for HC (0.26, 0.42, and 1.64) was descriptively larger than for GD (0.19, 0.22, and 1.13) (**Fig. S3**), with only sensitivity to loss being significantly larger in HC than in GD ($p_{WaldApprox} = 0.011$). LA was significantly smaller in GD than in HC ($p_{perm} < 0.001$).

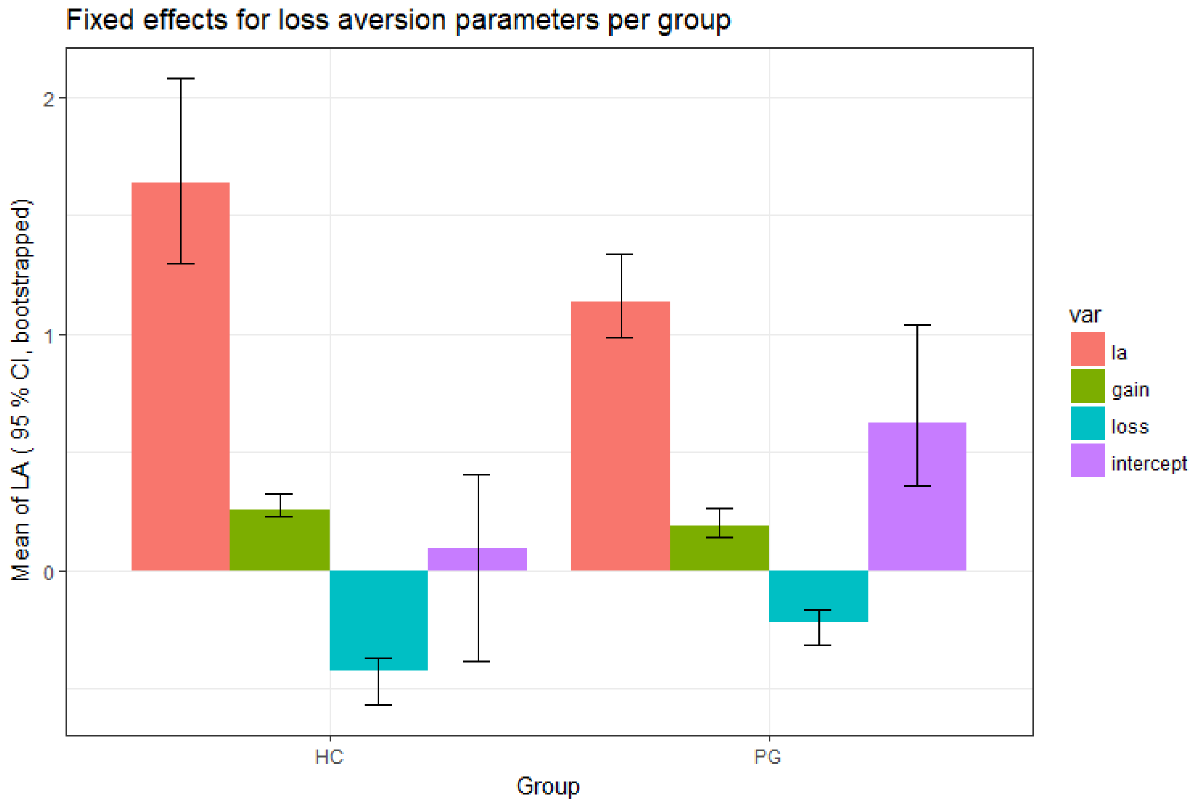


**Figure S3: Fixed effects loss aversion, gain sensitivity, loss sensitivity, and intercept per group.** The fixed effects and their non-parametrically bootstrapped CIs (many repetitions of lme fits with resampled subjects within groups) are displayed. var: variable, la: loss aversion, gain: gain sensitivity, loss: loss sensitivity, intercept: intercept of the logistic regression, i.e. the general acceptance rate at mean gain and loss

Adding the simple effect of category with group interaction lead to a significant improvement of the model ($p < 0.001$, $\Delta AIC = 691$). Here, we saw a significantly higher acceptance during gambling cues for GD subjects compared to HC ($p_{WaldApprox} < 0.001$). The additional triple-interaction "group X (gain, loss) X category" did not improve the model ($p = 1$, $\Delta AIC = -196$).

#### 2.3 Additional results graphs classifier

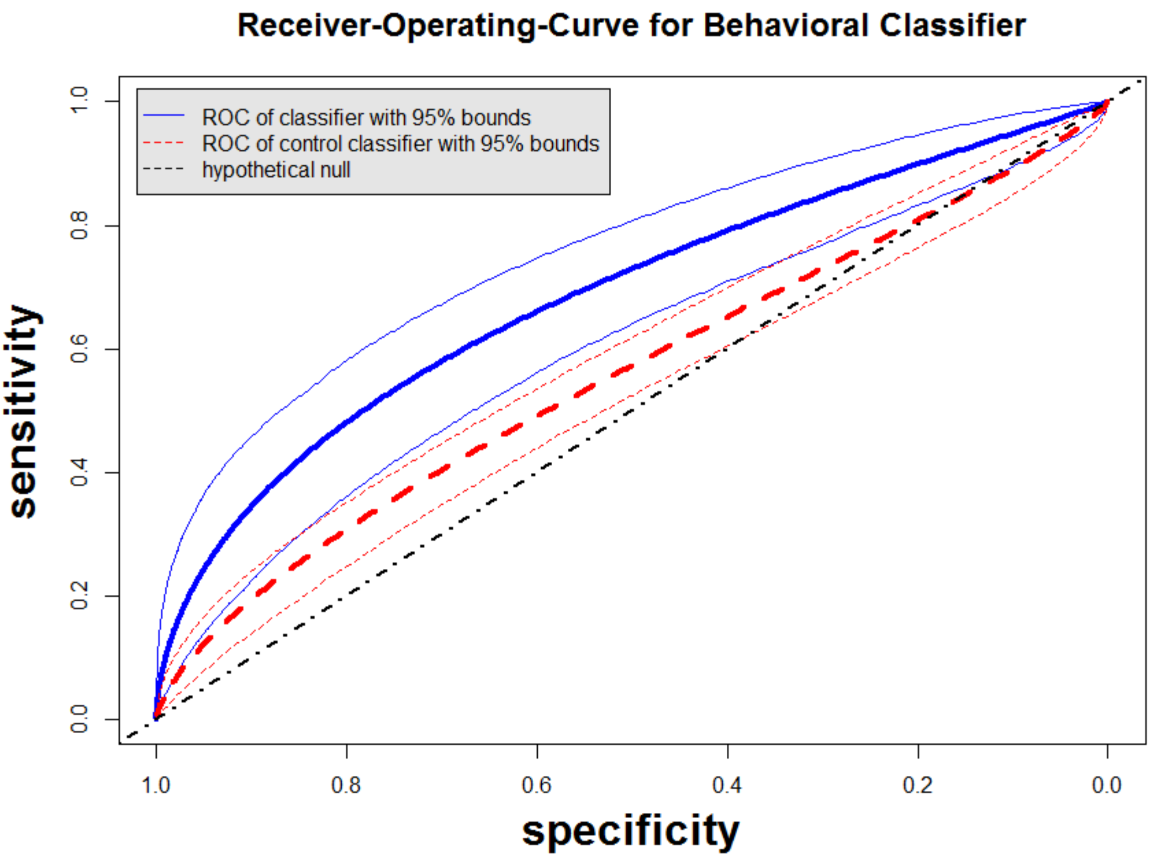


**Figure S4: Mean receiver-operating curve for behavioral classifier.** Describes how classifier fares concerning sensitivity and specificity when asked to distinguish GD subjects from HC subjects in independent subjects. Blue shows the ROC for the classifier (mean over 1000 rounds of the algorithm). Red is the ROC of the CV scheme with only smoking as predictor (0-hypothesis). Black is the theoretical 0-hypothesis. The classifier fares better than expected under the 0-hypothesis.

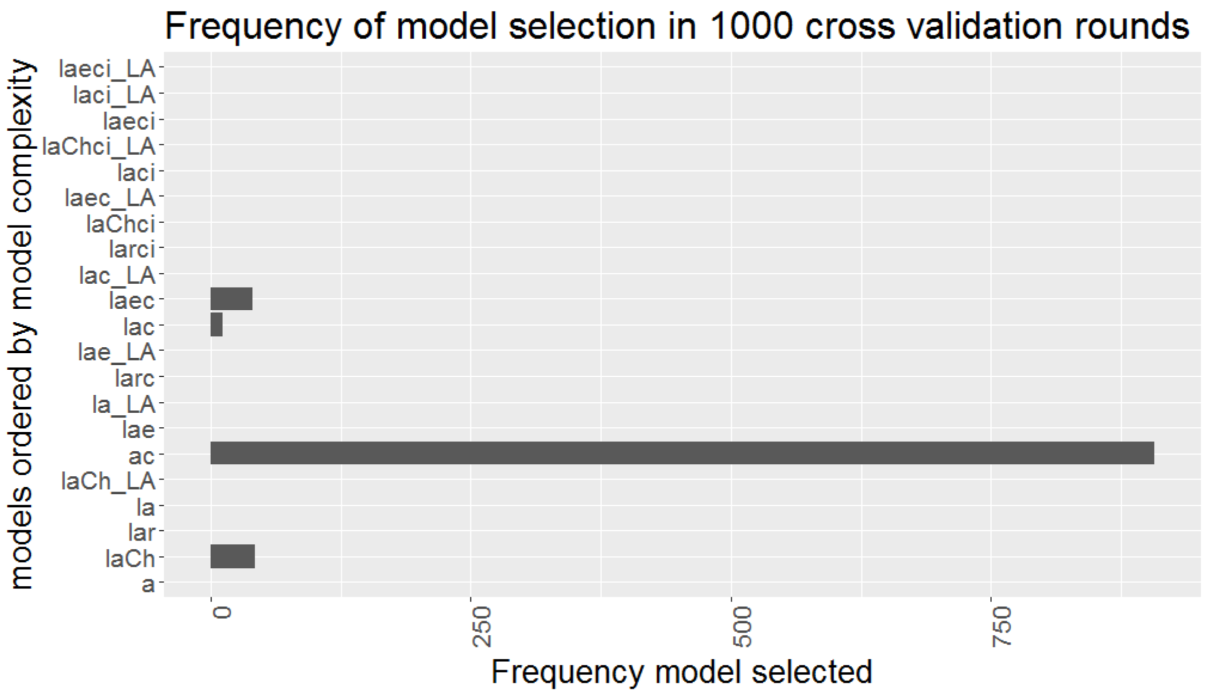


**Figure S5: Frequency of behavioral models selected.** The classification algorithm chose the “acceptance rate by category” model (**ac**) the most often in 1000 rounds of running the classification algorithm on the complete data. Models ending in “LA” are the models which have loss aversion (λ) parameters appended to the parameter set.

#### 2.4 Classifier results adjusted for cue repetition and with equal number of trials

We ran the algorithm by adding a factor "repeated_vs_novel" in each single-subject model as a covariate of no-interest, in order to adjust the estimation of all other parameters for that factor. In each trial, it was 0, if image was shown for the first time, else 1. We extracted all parameters per model and subject, as before, adjusted for "repeated_vs_novel". We also randomly selected 45 out of 67 gambling images to equalize the number of trials per cue category (now 45 in all categories). The results did not change meaningfully: AUC = 65.1, p = 0.016. On the validation set: AUC = 67.5, p = 0.017. The most often-picked model was still **ac** and its regression weights looked as before (**Fig. 2**). However, the selection of models was more varied now (**Fig. S6** compared to **Fig. S5**). When only adjusting for novelty and not cutting the gambling stimuli to 45, then the results were again like the original ones, and the distribution of selected models was very similar to **Fig. S5** (AUC: 64.5, p = 0.021, AUC on validation sample: 66.5%, p = 0.025. Thus, the cutting from 67 to 45 seems to make the difference in model selection, perhaps because 67 gamble trials just lead to stronger signal in the cue-dependent signal and thus the classifier uses also other models from time to time. However, note that still almost all models selected include the "c", i.e. the influence of category plays a role everywhere (on acceptance rate), just in some more models gain and loss sensitivity and loss aversion play a role additionally. In all analyses, cues have no relevant influence on gain and loss sensitivity (no interaction effect, **laci** model was not picked).

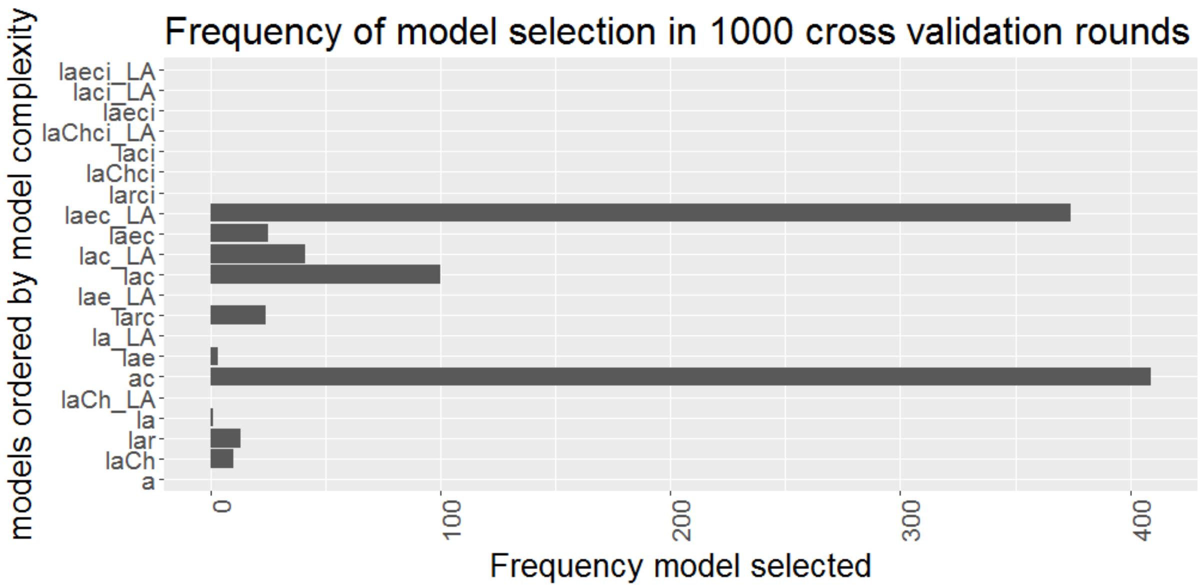


**Figure S6: Frequency of model selection in 1000 rounds of applying the algorithm.** Using only 45 gambling cues and adjusting for repeated presentation of cues.

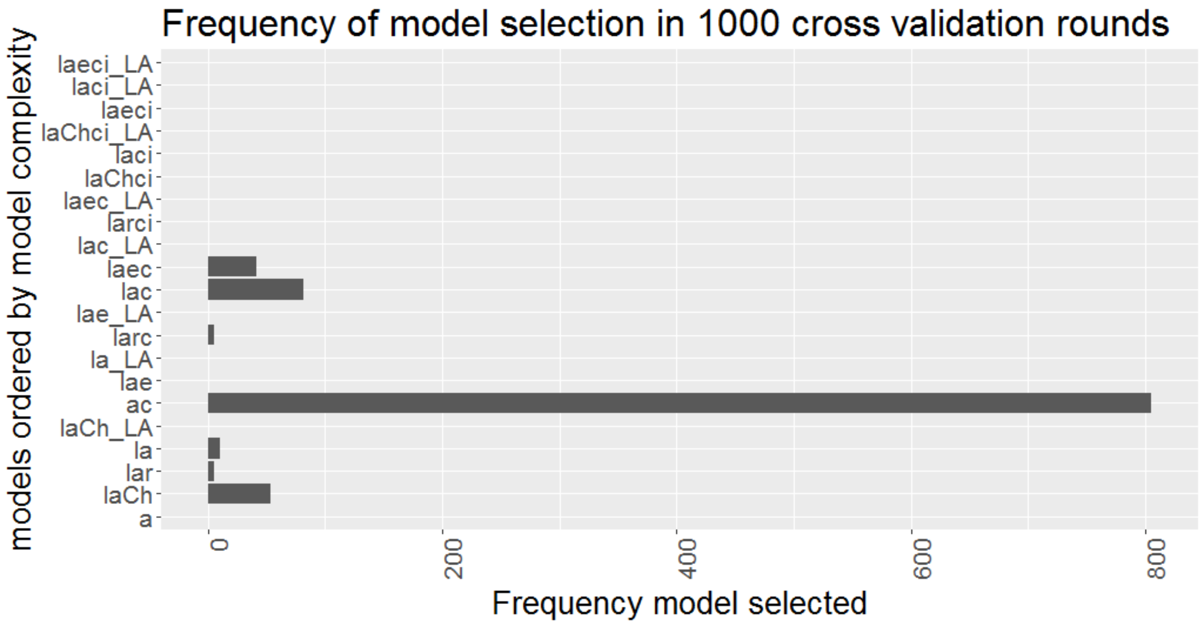


**Figure S7: Frequency of model selection in 1000 rounds of applying the algorithm.** Single-subject models adjusted for repeated vs. novel cues.
